# A hypothalamic-brainstem activity sequence underlies arousal fluctuations during daytime drowsiness

**DOI:** 10.64898/2026.05.26.728061

**Authors:** Ewa Beldzik, Nicholas G. Cicero, Daniel E.P. Gomez, Zinong Yang, Juan Eugenio Iglesias, Brian Edlow, Laura D. Lewis

## Abstract

During monotonous tasks, struggling with drowsiness leads to dynamic fluctuations in vigilance and task performance. Whether these moment-to-moment fluctuations are regulated by sleep–wake control circuitry remains unknown. Here, we used high-resolution 7T fMRI optimized for subcortical imaging to resolve activity across the human arousal regulatory system, including the hypothalamus, brainstem nuclei, basal forebrain, and thalamus, during a sustained attention task as subjects spontaneously experienced drowsiness. We identified a coordinated activity pattern locked to behavioral failures: widespread transient suppression during arousal drops and reactivation during recovery across all arousal-promoting regions, with opposite responses in the sleep-promoting hypothalamic preoptic area. These changes unfolded over ∼10–15 s in a 30 consistent sequence, led by the hypothalamic nuclei. Furthermore, suppression of brainstem nuclei preceded cortical infraslow (<0.05 Hz) oscillations, implicating a role in global neurovascular dynamics. These findings provide a temporally resolved, network-level portrait of human subcortical dynamics during drowsiness and identify a hypothalamic-led, brainstem-mediated activity sequence underlying behavioral arousal fluctuations and global cortical hemodynamics.

## Introduction

Sleepiness is described as a biological drive to sleep, yet it is not a simple transitional state preceding nighttime sleep. It is often experienced during the daytime without an obvious cause, and resisting it can be challenging even when motivation is high. Particularly during monotonous tasks, sleepiness intrudes in the form of sudden episodes resembling brief sleep and can resolve just as rapidly, resulting in pronounced fluctuations in cognition and behavior. Together, these observations suggest that daytime sleepiness is a dynamic phenomenon that may reflect a shifting interplay of sleep- and wake-promoting brain circuits. However, the neural circuit dynamics underlying daytime sleepiness, and in particular the brief drowsy episodes that disrupt task performance, are not well understood.

Pioneering rodent studies have delineated key subcortical arousal regulatory circuits, identifying a set of brainstem and hypothalamic nuclei that can causally induce sleep or wake states^1–6^. The system comprises morphologically heterogeneous nuclei in the brainstem, which release diverse neuromodulators, including the predominantly noradrenergic locus coeruleus (LC), the serotonergic median raphe (MnR) and dorsal raphe (DR), dopaminergic ventral tegmental area (VTA), cholinergic pedunculopontine tegmental nucleus (PTg), and glutamatergic-dominant regions such as the pontine reticular nucleus oralis (PnO), midbrain reticular formation (mRt), and periaqueductal gray (PAG)^7,8^. These brainstem nuclei and their axonal projections promote wakefulness by activating the thalamus and basal forebrain (BF), which together modulate arousal, consciousness, and attention^5,6,9, 10^. Conversely, hypothalamic nuclei regulate and stabilize arousal circuits, as proposed in the “flip-flop switch” model^11,12^. The preoptic area (POA) promotes sleep via GABAergic inhibition of arousal brainstem nuclei (OFF switch), while orexinergic neurons in the lateral hypothalamus (LH) stabilize wakefulness (ON switch). The tuberomammillary nucleus (TMN), the main source of histamine, promotes cortical arousal and inhibits the POA to favor wake states^13^. This fundamental knowledge about the mechanisms of sleep-wake regulation has been primarily derived from rodent studies — nocturnal species with fragmented sleep bouts — and remains scarcely explored in humans. Whether and how these interconnected subcortical circuits collectively operate to regulate the delicate balance between alertness and drowsiness during waking behavior remains unclear.

While the subcortical circuits underlying arousal fluctuations have not been extensively studied in humans, primarily due to the technical challenges of measuring activity in these small structures, many studies have identified profound changes in cortical activity during drowsiness. Sleep and sleepiness are characterized by the emergence of global cortical infraslow (<0.1 Hz) oscillations in the blood oxygenation level dependent (BOLD) signal^14–16^. In wakefulness, these oscillations are linked both to spontaneous arousal fluctuations in the resting-state^17,18^ and to microsleep intrusions during sustained attention^19,20^. Although the mechanisms driving these slow hemodynamic rhythms and their functional relevance remain poorly understood, they are tightly coupled to widespread brain-body physiological dynamics^21,22^, suggesting that they could reflect neuromodulatory state transitions originating within the arousal regulatory system^17,23^. Supporting this idea, various monoamines—including noradrenaline, dopamine, serotonin, and histamine—exhibit infraslow oscillations in sleep^24–27^ and have state-dependent vasoactive effects^28–30^, meaning that neuromodulatory activity of the arousal system could modulate global vascular tone and give rise to widespread cortical infraslow hemodynamics. However, whether specific subcortical activity dynamics underlie these drowsiness-linked cortical oscillations is not yet established.

Here, we imaged human subcortical arousal circuits during daytime drowsiness intrusions into an attentional task, aiming to identify the subcortical circuit dynamics underlying fluctuations in behavior, arousal, and cortical physiology. Functional imaging of these structures is technically challenging due to their small size and deep location in the brain^31^. We used an ultra–high-field (7 Tesla) fMRI protocol and analysis pipeline optimized for the subcortex, enabling high-resolution functional imaging with a high signal-to-noise ratio (SNR) across the entire network. We aimed to characterize how these subcortical arousal circuits dynamically interact in moment-to-moment fluctuations in alertness, by testing whether transient periods of unresponsiveness reflect coordinated network activity. Our second aim was to determine whether this subcortical circuitry contributes to global cortical infraslow BOLD oscillations, which would be expected if there is a neuromodulatory influence on drowsiness-related vascular physiology. We hypothesized that activity of the arousal regulatory system underlies both behaviorally defined arousal fluctuations and cortical vascular dynamics, thereby explaining the coupling between behavioral state and hemodynamics.

## Results

We imaged 27 healthy participants (mean age: 28.6±7.5 years; 11 males) during a sustained attention task. To increase drowsiness episodes, subjects slept only four hours the night before the scan, and scans were conducted in the early afternoon. To measure activity throughout the arousal system with high spatial precision, we designed a high-resolution (1.2 mm isotropic) 7T fMRI protocol optimized for the brainstem (Fig. 1A), acquired using a 64-channel coil^32^ designed for good subcortical coverage. Subjects performed the psychomotor vigilance task (PVT), a gold-standard task to quantify the effects of sleep deprivation^33^, as it requires sustained attention across long, randomized inter-trial intervals (Fig. 1B), which is impaired by increased homeostatic sleep pressure. Despite repeated instructions to stay alert and respond to every stimulus quickly and accurately, participants exhibited omission errors (failures to respond) on 20.5% (s.d = 0.4%) of trials, as expected from the sleep restriction paradigm. Two subjects who made fewer than two omission errors were excluded, resulting in a final sample of 25 participants (n = 25).

**Figure 1.**
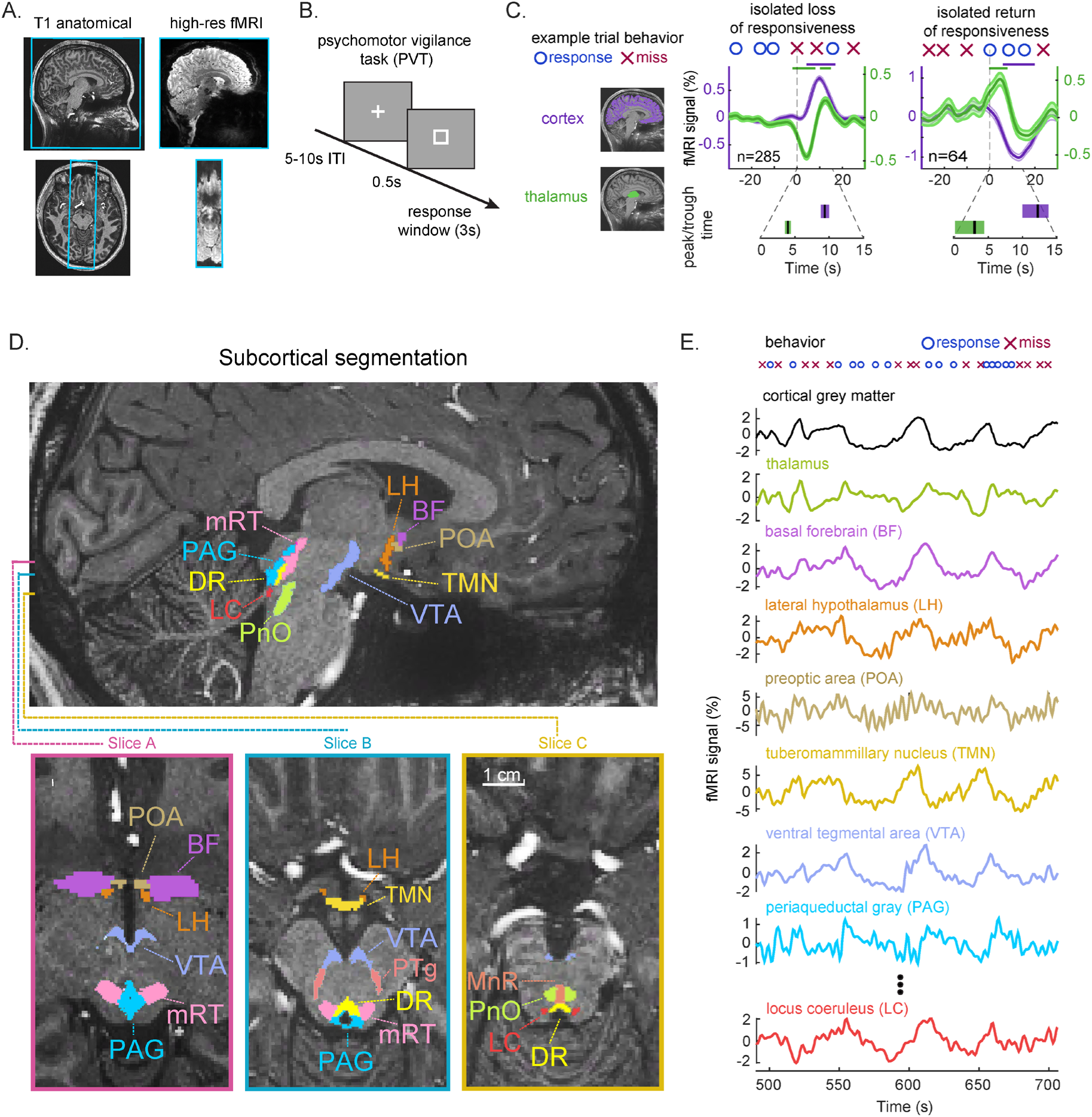
Experimental design and data analysis approach. (A) MRI protocol. High-resolution (1.2 mm isotropic) gradient-echo EPI was acquired sagittally (blue outline) to cover the brainstem, midbrain, thalamus, and midline cortical regions. The anatomical segmentations were linearly registered to distortion-corrected functional space. Functional images therefore remained in their original space, to minimize spatial transformations that could compromise the signal in small subcortical regions. (B) Experimental task. Subjects performed the psychomotor vigilance task (PVT) by pressing a button in response to every square stimulus as rapidly and accurately as possible. (C) Mean global cortical (purple) and thalamic (green) fMRI signals, locked to isolated arousal transition events. Signals locked to the first omission trials (loss of responsiveness, left panel) and correct trials (return of responsiveness, right panel) showed a sleep-like global cortical hemodynamic response with a delayed and inverted response relative to subcortical brain activity. Top panel: Colored shading indicates standard error of the mean. Horizontal lines indicate significant differences (FDR-corrected p<0.05, linear mixed-effect with random intercept for subject) from the baseline (-30 s to -20 s). Bottom panel: mean peak/trough timing relative to trial onset. Color shading is 95% confidence intervals. (D) Segmentation of subcortical ROIs in an example subject’s anatomical space. (E) Example fMRI timeseries in cortical gray matter, thalamus, hypothalamus, and selected brainstem regions showing strong amplitude changes linked to behavioral responses (top plot).

### Sustained attention is disrupted by microsleep-like behavioral lapses

Such profound behavioral impairment on a simple, low-effort task suggests that participants experienced intrusions of microsleep episodes^19,20^, rather than mere reductions in vigilant attention^34^. To examine this possibility, we first aimed to test whether these periods of unresponsiveness resembled microsleeps, or sleep intrusions, based on the hemodynamic response profiles in the cortex and thalamus. Specifically, previous studies have shown that sleep onset is locked to thalamic suppression followed by a rise in the cortical signal^19^, whereas awakening from sleep is characterized by the opposite sequence—namely, a thalamic increase followed by cortical suppression^20,35^. Similar to Setzer and colleagues (2022), we extracted fMRI signals from thalamic and cortical regions and time-locked them to moments of a behavioral state change after a long stable period. Specifically, we examined omission trials preceded by at least 30 s of correct responses (isolated loss of responsiveness, n = 285), and correct trials preceded by at least 30 s of omissions (isolated return of responsiveness, n = 64). In line with previous results, we found that the subcortical and cortical patterns linked to entering and recovering from these periods of unresponsiveness replicated the patterns known to occur at sleep onset (Fig. 1C), consistent with microsleep-like intrusions into wakefulness.

### Arousal transitions are locked to coordinated suppression and activation of subcortical arousal regions

To identify subcortical regions and nuclei involved in arousal transitions, we applied recently developed segmentation tools^8,36–38^ that enable precise localization in each participant’s native anatomical space using T1-weighted images (see example segmentation in Fig. 1D), avoiding spatial blurring from group atlases that compromises signal fidelity in these small structures. Our regions of interest (ROIs) encompassed eight brainstem nuclei (LC, MnR, DR, VTA, PnO, mRt, PAG, PTg), three hypothalamic subregions (LH, POA, TMN), as well as the BF and thalamus—key brain regions of the sleep–wake regulatory circuitry according to animal studies^5,6^. We registered these ROIs to the preprocessed, distortion-corrected functional images and extracted fMRI timeseries from each ROI (see example timeseries in Fig. 1E).

We first examined activity within the sleep–wake regulatory circuitry during transitions into and out of drowsiness by identifying behaviorally defined changes in arousal for each subcortical ROI. In contrast to the analysis in Fig. 1, here we included all trials representing arousal transitions; specifically, omission trials preceded by at least one correct trial (loss-of-responsiveness, n = 955 trials) and correct trials preceded by at least one omission trial (return-of-responsiveness, n = 936 trials). Although this approach introduces exposure to the effects of preceding arousal events due to temporal overlap between consecutive transitions, it includes all drowsy events (which can be brief). We avoided the influence of preceding events by restricting analyses to the −5 s to +30 s window relative to event onset, a period we verified to be unaffected by these prior arousal events in the subcortex (Extended Data Fig. 1).

Consistent with their established wake-promoting roles^5,9,12^, TMN, LH, all brainstem nuclei, as well as thalamus and BF exhibited decreased activity during loss of responsiveness and increased activity during recovery (Fig. 2A). In contrast, and in line with its sleep-promoting role^12,39^, the POA was the only region showing the opposite pattern, with activation prior to drowsiness onset, and deactivation prior to recovery of responsiveness. The peak and trough amplitudes of hypothalamic activity occurred slightly before time zero, corresponding to onset of the behavioral change. Accounting for the ∼5 s delay introduced by the hemodynamic response in fMRI signals relative to underlying neural activity, these findings suggest that arousal-network dynamics begin to appear several seconds before the system transitions out of or back into responsiveness. Together, the results are consistent with a neuromodulatory state control framework^23,40^, in which tonic arousal system activity regulates cortical and subcortical excitability, thereby setting the threshold for maintaining behavioral responsiveness under sleep restriction. Furthermore, our results suggest that this regulatory process is relatively slow, with coordinated activity across regions unfolding over multiple seconds prior to behavioral state changes.

**Figure 2.**
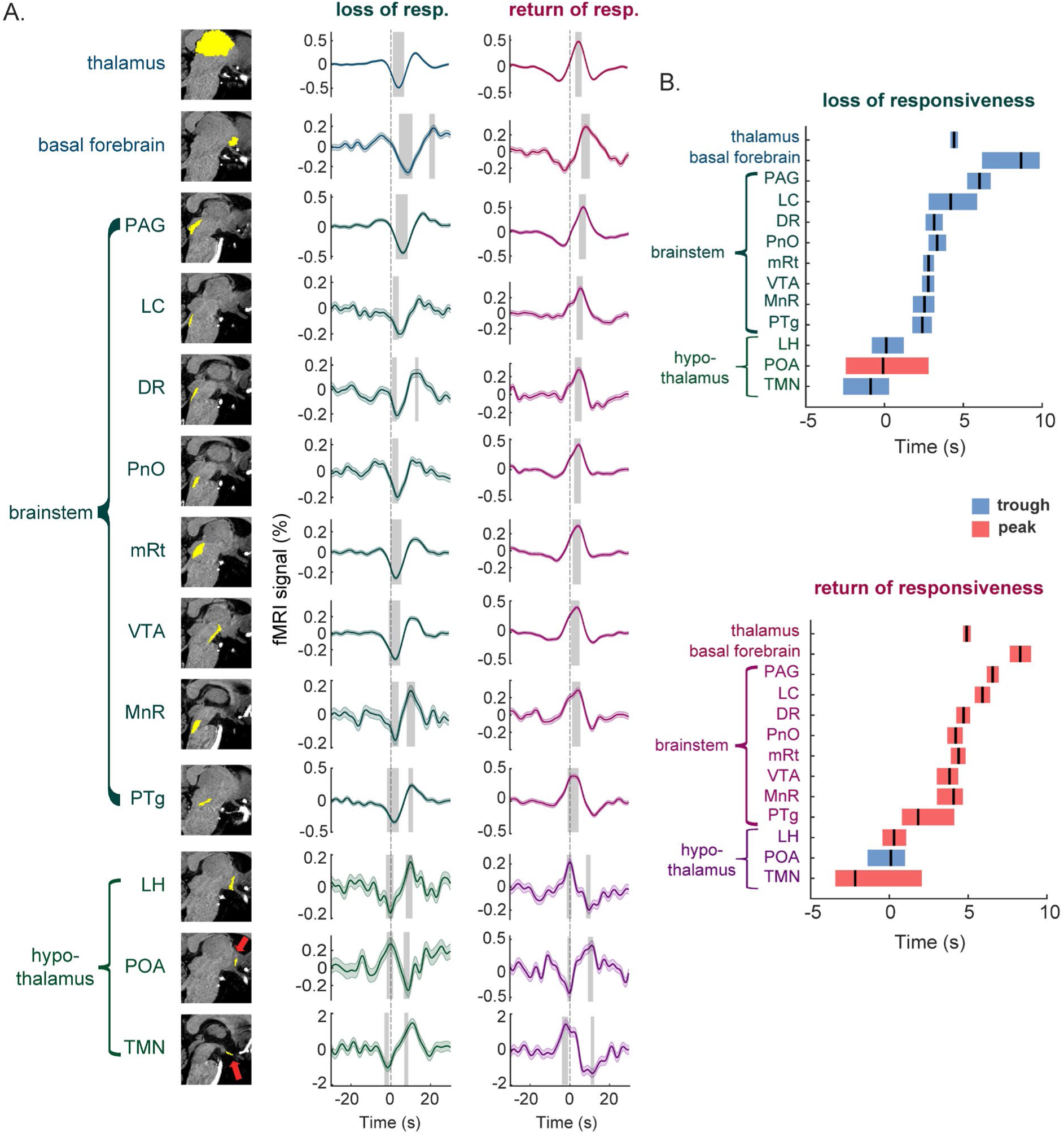
Coordinated activity across subcortical nuclei is locked to transitions in behavioral arousal state. (A) Mean fMRI signal in each ROI, locked to the onset of omission trials (loss of responsiveness, green) and the onset of correct trials (return of responsiveness, purple). Colored shading indicates standard error of the mean. Grey shading indicates significant changes (FDR-corrected p<0.05, linear mixed-effect with random intercept for subject) from baseline (-30s to -20s). All regions show significant activity locked to arousal state changes; the direction of this change is opposite in POA relative to other regions. N=955 ‘loss’ trials, 936 ‘return’ trials. (B) Average timing of the first significant peak or trough in each region’s trial-locked response. Color shading indicates 95% confidence intervals (bootstrap). PAG=periaqueductal gray, LC=locus coeruleus; DR=dorsal raphe, PnO=pontine reticular nucleus oralis, mRt=midbrain reticular formation, VTA=ventral tegmental area, MnR=median raphe; PTg=pedunculopontine tegmental nucleus, LH=lateral hypothalamus, POA=preoptic area, TMN=tuberomammillary nucleus.

### Hypothalamic nuclei lead brainstem, thalamic, and basal forebrain activity during behavioral state changes

We next asked whether activity changes across the arousal regulatory regions occurred synchronously or whether specific regions led the behavioral transitions. To quantify response timing, we estimated the latency to the first prominent peak or trough in each condition and derived 95% confidence intervals using trial-wise bootstrapping. We found that activity changes in hypothalamic nuclei preceded those in the brainstem and thalamus by approximately 3–8 s, depending on the region (Extended Data Table 1), and were followed by delayed activity in the basal forebrain in both conditions (Fig. 2B). While differences in neurovascular coupling timing can contribute to apparent delays in fMRI signals across regions^41–44^, those differences are typically <2 s, substantially smaller than the differences across ROIs observed here. Therefore, the magnitude and the consistent ordering of these latencies, together with the opposite response profile observed in the POA, argue against explanations based solely on differences in vascular anatomy across ROIs and instead support a sequential engagement of arousal circuitry during behavioral state transitions.

### Global cortical infraslow BOLD oscillations are preceded by suppression of brainstem, thalamic, and basal forebrain activity

Given that this subcortical activity sequence preceded the behavioral effects of drowsiness, we next investigated whether this activity also serves as a potential neuromodulatory basis of global cortical infraslow oscillations, a key cortical signature of arousal state in humans. We first confirmed that drowsiness, defined as the omission rate within a 1-min window, was associated with increased BOLD signal power, most prominently in the infraslow (0.01–0.05 Hz) range (Extended Data Fig. 2). We filtered the global cortical fMRI signal within this frequency range and detected prominent (>1 s.d.) peaks across all runs (Fig. 3A). In line with prior work^22^, cortical peaks (n = 1274) were preceded by a sharp increase in omission likelihood (Fig. 3B), suggesting that transient microsleep intrusions may evoke this sleep-like cortical vascular response.

**Figure 3.**
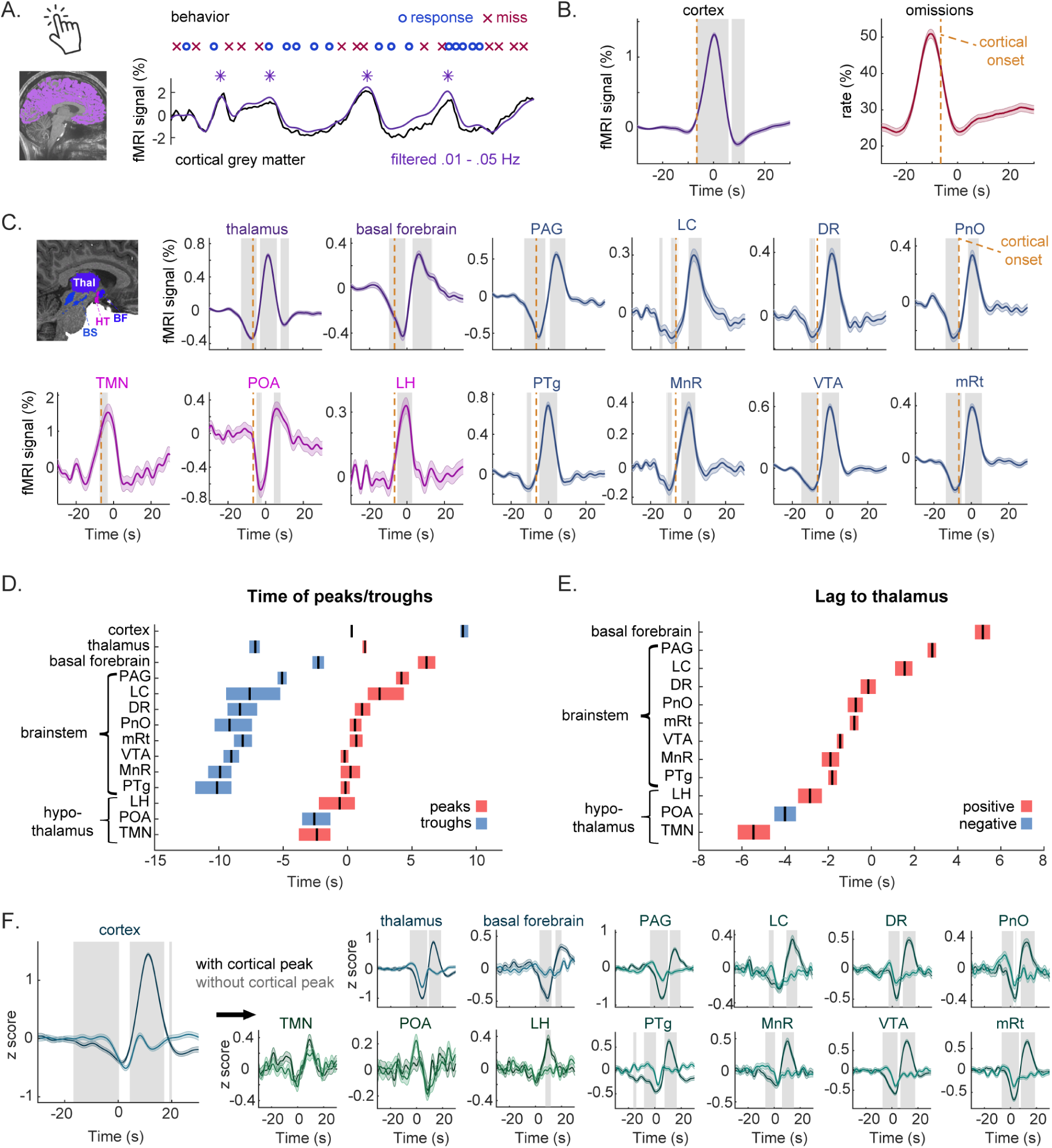
A structured, biphasic subcortical activity sequence precedes the global cortical peaks of infraslow fMRI oscillations. (A) To identify cortical peaks, the fMRI signal in cortical gray matter was filtered in the 0.01–0.05 Hz range, and peaks exceeding 1 standard deviations were extracted. (B) The average cortical fMRI signal locked to infraslow cortical peaks (left). Peaks are preceded by a robust increase in the rate of behavioral omissions (right). Color shading indicates standard error of the mean. Gray shading indicates significant changes (FDR-corrected p<0.05, hierarchical bootstrapping to account for subject-level variability) from baseline (–30 s to –20 s). Dashed orange line marks the onset of significant change in cortical signal amplitude. (C) fMRI signal in the arousal regulatory system locked to global cortical peaks. Color shading indicates standard error of the mean. Gray shading indicates significant changes (FDR-corrected p<0.05, hierarchical bootstrapping to account for subject-level variability) from baseline (–30 s to –20 s). All regions, except hypothalamus, showed decreased activity preceding the onset of cortical rise. (D) Averaged timing of the significant peaks or troughs relative to cortical peak (time zero). Color shading indicates 95% confidence intervals. (E) Mean time lag of the regions relative to the thalamus. Color shading is 95% confidence intervals. (F) Averaged fMRI signal in arousal regions in the loss-of-responsiveness condition with and without a following global cortical rise. Color shading indicates standard error of the mean. Gray shading indicates significant differences (FDR-corrected p<0.01, linear mixed-effect with random intercept for subject) between the peak and no-peak conditions. HT=hypothalamus, PAG=periaqueductal gray, LC=locus coeruleus; DR=dorsal raphe, PnO=pontine reticular nucleus oralis, mRt=midbrain reticular formation, VTA=ventral tegmental area, MnR=median raphe; PTg=pedunculopontine tegmental nucleus, LH=lateral hypothalamus, POA=preoptic area, TMN=tuberomammillary nucleus.

We next analyzed fMRI timeseries across all subcortical regions, locked to the peaks of cortical infraslow oscillations, to examine whether their activity temporally preceded the cortical wave onset, defined as the first timepoint with a significant increase from baseline. This analysis revealed a significant decrease in BOLD signal within the brainstem, thalamus, and BF preceding the onset of cortical waves (Fig. 3C). The temporal profile of these deactivations indicated that PTg and MnR significantly preceded thalamus, PAG, and BF (Fig. 3D; Extended Data Table 2). Notably, the relatively late BF deactivation, occurring nearly in parallel with the cortical peak, is consistent with prior work showing an anticorrelation between BF activity and global cortical peaks in the resting state^17^. These results demonstrated that this cortical–BF relationship was preceded by a coordinated cascade of deactivation across arousal regulatory nuclei, together with declining vigilance, with BF activity changes as the final step in this sequence. Overall, these results identify a structured sequence of subcortical activity that consistently precedes the induction of a cortical infraslow wave.

Cortical peaks were linked to a biphasic change in behavior: a strong increase in lapses at -10s, followed by a subsequent recovery of behavior. Parallel to this behavioral profile, the subcortical activity patterns were also clearly biphasic, with decreases before the cortical peak, and increases before the subsequent cortical trough, consistent with the reversed and delayed pattern of cortical activity observed during sleep–wake transitions^20,35^. The subcortical activation pattern linked to behavioral recovery was led by hypothalamic nuclei, followed by the brainstem, thalamus, and, lastly, the BF, consistent with the behavior-based return-of-responsiveness analysis. To enhance sensitivity in distinguishing the temporal characteristics of these signals across regions, we performed a trial-by-trial time-lag analysis between the thalamus and the remaining arousal-regulatory regions. This analysis demonstrated a clear temporal cascade of activation, with the hypothalamic TMN region leading these recovery-related fMRI signal changes (Fig. 3E; Extended Data Table 2).

### Subcortical arousal dynamics independently explain behavioral lapses and cortical infraslow activity

Together, these findings indicate that arousal system suppression, and POA activation, underlies drowsiness-related processes at two levels: behavioral (periods of unresponsiveness consistent with microsleeps) and physiological (enhanced global cortical infraslow rhythms). These processes are tightly coupled, likely reflecting a shared neuromodulatory influence on neural activity and vascular physiology. We next asked whether the network suppression contributes independently to behavioral decline and the subsequent cortical peak.

To test this, we reexamined the loss-of-responsiveness condition (Fig. 2, green traces) by comparing omission trials with and without a subsequent cortical peak. We expected that omission trials followed by a pronounced global cortical hemodynamic response would reflect greater suppression of the arousal system than omission trials with limited cortical changes. We therefore divided trials based on the median amplitude of the average cortical BOLD peak (∼11 s after omission onset) and averaged the fMRI time series for each subcortical region in these two conditions (peak and no-peak). Trough amplitudes in the thalamus, BF, and all brainstem nuclei were significantly larger in trials associated with stronger subsequent cortical peaks (Fig. 3F), indicating additional suppression of these regions during arousal dips accompanied by a vascular response. These results confirm that subcortical circuitry involving the brainstem, thalamus, and BF jointly underlies both the behavioral decline and the global vascular response during drowsiness.

Notably, hypothalamic regions did not show activity linked to the cortical oscillation peak amplitude: they showed neither a pre-cortical decrease (Fig. 3C) nor differences between trials with or without a subsequent cortical peak (Fig. 3F). This pattern suggests either a limited direct contribution of hypothalamic nuclei to the generation of cortical infraslow rhythms, or an indirect involvement mediated by slower or more tonic neuromodulatory processes not captured by the present analysis. To further examine this possibility, we applied a linear mixed-effects model to behavior-locked data, in which behavior-related activity was captured in the intercept term, while variability in cortical peak amplitude was included as a separate regressor (Extended Data Fig. 3). This analysis revealed subtle modulation of hypothalamic activity by cortical amplitude, consistent with mild and slow influences on cortical hemodynamics. Interestingly, parameter estimates associated with cortical peak amplitude in all regions reached their maxima earlier than omission-related effects, suggesting that neuromodulatory influences on the cortical infraslow oscillation, across the arousal system, are temporally dissociable from behavioral arousal transitions.

## Discussion

This study identified a hypothalamic-brainstem activity sequence that underlies fluctuations in sustained attention performance when struggling with daytime drowsiness. We observed distinct temporal dynamics across arousal regulatory circuits during sleepiness-induced arousal transitions, coupled both to behavioral loss and return of responsiveness, and to infraslow cortical oscillations. The activity patterns in subcortex were highly local, in contrast to the large, global cortical dynamics during unintended intrusions of sleepiness during sustained attention. We further found that the sequence of hemodynamic responses across the arousal circuitry unfolds over a prolonged temporal profile of approximately 10–15 s. Notably, hypothalamic nuclei were the earliest to show arousal-related activity before behavioral changes, suggesting that maintaining alertness following sleep restriction depends on the stabilizing role of this region, in line with the canonical flip–flop model of sleep–wake regulation developed in rodents^11,12^. Together, our results provide a comprehensive depiction of how activity unfolds throughout arousal circuits in humans during drowsy episodes, and reveal a characteristic timescale governing these processes.

The early and robust engagement of the TMN suggests that this region may play a critical role in resisting sleep pressure. Previous animal studies have demonstrated a central role for histaminergic neurotransmission in stabilizing arousal^13,45^, acting either indirectly via the POA, LC, raphe nuclei, thalamus, and BF^46,47^, or directly through projections to the cortex and striatum^48^. Our findings extend this work by providing system-level evidence in humans that this activity is tightly temporally linked to behavioral task performance. This imaging approach could open new avenues for investigating the mechanisms underlying the effects of widely used over-the-counter sedatives, such as H₁-antagonist antihistamines, as well as recently approved wake-promoting treatments for narcolepsy based on histamine H₃ receptor inverse agonists^49^.

The hypothalamic POA exhibited similarly early timing to the TMN, significantly preceding brainstem nuclei, but with opposite activity during arousal transitions, demonstrating that our high-resolution imaging approach can detect focal activity in this local sleep-promoting nucleus. This ability to image POA in humans could next be used to investigate sleep disorders, such as excessive daytime sleepiness. The POA exerts predominantly inhibitory, GABAergic and galaninergic efferent projections to arousal-promoting regions, particularly the TMN, LH, and several monoaminergic brainstem nuclei^50^. In turn, the POA is inhibited by afferent monoaminergic inputs from these regions, positioning it as a central hub for bidirectional regulation of sleep and wakefulness^39^. Despite this well-established circuitry, little is known about POA involvement in attentional failures during sustained, self-motivated attention. Our findings suggest that microsleep intrusions into wakefulness may be initiated by POA activation and suppression of the TMN and LH, leading to subsequent suppression of monoaminergic brainstem nuclei and the thalamus. By contrast, recovery from these drowsy periods is initiated by TMN activation and POA inhibition, leading to sequential excitation of brainstem nuclei, starting with the cholinergic PTg.

We next captured arousal fluctuations during infraslow cortical BOLD oscillations, which—given their global cortical distribution—are expected to reflect changes in vascular tone and autonomic state^51,52^, in addition to large-scale neuronal activity. The norepinephrine–LC system has been identified in animal models as a dominant driver of systemic physiological arousal responses^53,54^ and a key regulator of cortical vasoconstriction during sleep^24,55^. In the present study, we observed transient deactivation not only within the LC, but across all brainstem nuclei preceding the global BOLD signal increase, consistent with recent findings that many arousal-related neuromodulatory systems exhibit synchronized infraslow cortical oscillations during NREM sleep^25,26^. Although this activity pattern was associated with behavioral unresponsiveness and followed a similar temporal sequence in behavior-locked data, the two phenomena could also be dissociated. Specifically, behavioral lapses did not necessarily coincide with the emergence of a cortical infraslow wave, suggesting that momentary failures to respond can sometimes occur without a global cortical state transition. A parsimonious interpretation of these results is that only sufficiently strong suppression of brainstem nuclei leads to inhibition of the norepinephrine–LC system, resulting in global cortical vasodilation and a BOLD signal increase. By contrast, weaker brainstem suppression still elicits robust drowsy episodes, visible as decreases in attentional performance. This dissociation may reflect mechanistic differences between local sleep–like processes or mind blanking^56–58^ and globally expressed microsleep states^19,20,59^, although direct validation will require simultaneous EEG recordings^60,61^. Notably, BF activity was more tightly coupled to the cortical infraslow oscillation than to behavior, implicating this region as a potential determinant of whether an arousal dip results in a transient decrease in vigilance or progresses to a global microsleep^9,62^.

Together, these observations demonstrate that transient activity within subcortical arousal circuits can be reliably measured using ultra–high-field 7T fMRI, providing a framework for studying the human arousal system and offering new insight into how subcortical circuits orchestrate transitions between wakefulness and sleepiness. By enabling precise mapping of neuromodulatory activity in vivo, our imaging approach opens new opportunities to investigate the pathophysiology of sleep disorders such as insomnia and hypersomnolence and to assess the systems-level effects of stimulants and hypnotic drugs. Future studies should also extend this framework by identifying the subcortical systems that selectively regulate autonomic signals coupled to global infraslow hemodynamic oscillations^18,22,52^. Our results reveal how coordinated neuromodulatory dynamics in the subcortex shape global brain states, offering a foundation for future studies probing pathological and pharmacological alterations of arousal.

## Materials and Methods

### Participants

Data were acquired from 27 healthy subjects (16 females, 11 males) with mean age of 28.6 ± 7.5 (std). All were right-handed although it was not an inclusion criterion. All experimental procedures were approved by the Boston University Charles River Campus Institutional Review Board (IRB 5059E). Exclusionary criteria included all MRI contraindications, as well as any sleep, neurological, or psychiatric disorders and related medications. All subjects provided written informed consent before participating in the study. Subjects reported good typical sleep quality measured with Pittsburgh Sleep Quality Index (PSQI) questionnaire with average score of 4.2 ± 2.1 (std) and normal levels of daytime sleepiness measured with Epworth Sleepiness Scale (ESS) with average score 6.2 ± 3.5 (std). Participants were instructed to restrict their sleep in the night prior to the scan to four hours total. Compliance with this instruction was verified with a wrist-worn actigraph (ActiGraph wGT3X-BT, ActiGraph, LLC) that was worn for 24 hours before the MRI session, and with sleep diaries filled out the morning of the scan session. Participants were asked to refrain from caffeine intake for 6 hours and alcohol for 24 hours prior to MRI session. Scans were scheduled at 1 pm or 3 pm to induce drowsiness due to the post-lunch dip^63^. As indicated by actigraphy (n=23), subjects slept 234 min ± 50 (std) the night before the scan. On the morning of the MRI visit, they reported subjective sleepiness of 3.0 ± 1.1 (std) measured with Stanford Sleepiness Scale (SSS). The MRI scan took place on average 7h and 23 min after awakening, measured by actigraphy.

### Experimental task

The PVT task was implemented using Psychtoolbox^64^ and projected onto a screen positioned at the end of the MRI bore, visible by mirror placed inside the MRI headcoil. Subjects were instructed to respond as fast and accurately as possible by pressing a button with the index finger of the dominant hand when the fixation crosshair changed into a square (Fig. 1B). The square was designed to have identical luminance to the fixation crosshair to avoid light-induced arousal responses. Each stimulus lasted 0.5s and inter-trial intervals ranged from 5-10 s. The stimuli were presented approximately 100 or 150 times during the 12-min or 18-min long task, respectively.

### MRI acquisition

Subjects were scanned on a Siemens 7T Terra scanner using a custom 64-channel head and neck coil^32^. The anatomical scan included a T1-weighted MPRAGE sequence (voxel size = 0.75 mm isotropic; 224 slices; TR = 3.82 s; TA = 7.53 min). Functional data (Fig. 1A) collected during the PVT task consisted of a high-spatial-resolution sagittal partial-volume gradient-echo EPI acquisition (voxel size = 1.2 mm isotropic; 38 slices; TR = 0.98 s; TE = 22.6 ms; SMS = 2; GRAPPA = 4; phase encoding = A–P). Sagittal acquisition volumes were positioned to provide coverage of the midline and brainstem. Each participant completed either four 12-min functional runs (735 volumes) or three 18-min runs (1102 volumes). Run durations were 12 minutes for subjects 1–11 and 18 minutes for subjects 12–27. Starting with subject 12, the scanning protocol additionally included reverse phase-encoding images matching the field of view of the functional scan (voxel size = 1.2 mm isotropic; TR = 0.98 s; TE = 22.6 ms; SMS = 1; GRAPPA = 4; 10 volumes), acquired separately for P–A and A–P encoding directions to enable distortion correction using TOPUP^65^. After each functional run, subjects completed the Stanford Sleepiness Scale (SSS), presented on the screen, indicating their current sleepiness level using a button box. Peripheral physiological monitoring was performed using a pulse oximeter, a piezoelectric respiratory belt, and galvanic skin response (GSR) sensors.

### MRI preprocessing

Anatomical MRI data were bias-field corrected using SPM12^66^, and segmented using FreeSurfer v7.4^67^. For subcortical segmentation, we used four recently developed segmentation tools that enable high-precision nucleus segmentation within individual subject space: 1) Brainstem Nuclei Segmentation Tool^8^ for all brainstem nuclei, 2) Hypothalamic Segmentation Tool^36^ for POA, within anterior-superior hypothalamus subunit, 3) FastSurfer-HypVINN, Hypothalamic Segmentation Tool^38^ for TMN, within tuberal region, and LH, 4) Subcortical Limbic Segmentation Tool for BF^37^. Notably, the thalamus was analyzed as a single region, as the temporal sequencing of activity across individual thalamic nuclei during arousal transitions has been characterized previously^35^.

Functional MRI data were first denoised using NORDIC^68^ to reduce contributions of thermal noise. The data were then slice-time corrected using FSL v5^69^ and motion corrected using AFNI v2016^70^. Next, reverse phase-encode (TOPUP) maps were applied to correct distortions in the A–P direction using the AFNI *3dQwarp* function. Datasets without TOPUP maps were corrected for distortion using nonlinear registration optimized over a subcortical mask implemented in ANTs^71^. All registrations were manually inspected to ensure accurate localization and to avoid partial voluming with adjacent spaces, such as the fourth ventricle (Fig. 1C). Spatial registration was conducted to ensure that functional images remained in their original space, minimizing transformations that could compromise signal quality in small subcortical regions. Thus, individual T1-weighted anatomical images for each subject were rigidly registered to distortion-corrected functional space using the FreeSurfer *bbregister* function^72^. The resulting transformation matrix was applied to all segmented regions of interest (ROIs), using a 0.7 threshold to fill the original anatomical masks. Any overlapping voxels between neighboring ROIs were excluded. Subjects with ROI functional masks comprising fewer than 2 voxels were excluded from further analysis. As a result, the POA and TMN regions were excluded in 2 and 10 subjects, respectively.

We took extensive measures to account for motion in our data. First, we applied FSL’s MELODIC^73^ decomposition separately to fMRI data masked for cortex, thalamus, hypothalamus, limbic regions, and brainstem, using an arbitrary but commonly used number of components (n = 20). The resulting component time series were correlated with the six motion time series. Any component with an absolute correlation > 0.7 with any motion trace was excluded from the subsequent reconstruction of the fMRI signal. On average, 3.9 ± 1.9 (SD) components were removed in each decomposition. Second, we interpolated the reconstructed fMRI time series at any time point for which motion RMS exceeded 2 standard deviations. This resulted in 3.7% ± 1.5% (SD) of data being interpolated. Data were then converted to percent signal change and filtered from 0.01 Hz to 0.2 Hz. Next, time series were interpolated to 0.01 s sampling rate using spline interpolation. All further data analyses were conducted only for those fMRI session runs in which subjects committed more than one omission error, ensuring a sufficient level of drowsiness and enabling omission-locked data analysis. Applying this criterion resulted in excluding 12 runs (out of 74). The final sample size in the study was 25 subjects, with an average of 2.5 functional runs per subject.

### Loss and return of responsiveness data analysis

We first aimed to investigate fMRI signals in global cortical grey matter and subcortical regions combined (i.e., a mask including all subcortical ROIs) during isolated events of loss and return of responsiveness. Specifically, we time-locked the fMRI timeseries extracted from these regions to the first omission trial preceded by at least 30 s of correctly responded trials (isolated loss-of-responsiveness condition, n = 285 trials), serving as a proxy for entry into (micro)sleep, or to the first correct trial preceded by at least 30 s of missed trials (isolated return-of-responsiveness condition, n = 64 trials), representing recovery from a drowsy state. These epoched data were baseline-corrected by subtracting the mean signal value between the −30 s to −20 s time range for each trial. Next, each time point was tested for significant change from baseline using a linear mixed-effects model according to the formula ‘fMRI amplitude ∼ 1 + (1 | subj)’ to account for subject-level variability. The obtained *p* values were corrected for multiple comparisons using the FDR method at the 0.05 level. We next estimated the latency of troughs and peaks for each ROI and behavioral condition using non-parametric bootstrapping by resampling trials with replacement (1,000 iterations), computing the mean timeseries, and extracting the largest negative and positive deflections within a −5 to +15 s window. The resulting bootstrap distributions of peak and trough latencies were used to calculate 95% confidence intervals (Fig. 1E).

To address the first goal of the study, that is, to identify transient hemodynamic activity in the sleep–wake regulatory circuitry during loss- and return-of-responsiveness conditions, we included all arousal events because (1) they represent meaningful events that characterize sustained attention impaired by drowsiness, and (2) they provide robust statistical power necessary to investigate regions with low signal-to-noise ratio. Specifically, we time-locked the fMRI timeseries extracted from each ROI to the first omission trial (loss-of-responsiveness condition, n = 955), and to the first correct trial (return-of-responsiveness condition, n = 936 trials), These epoched data were baseline-corrected by subtracting the mean signal value between the −30 s to −20 s time range for each trial. Next, each time point was tested for significant change from baseline using a linear mixed-effects model according to the formula ‘fMRI amplitude ∼ 1 + (1 | subj)’ to account for subject-level variability. Statistical testing was limited to −5 s to +20 s window range to avoid potential effects of the previous arousal event (Extended Data Fig. 1). The obtained *p* values underwent multiple-comparison correction using the FDR method at the 0.05 level (Fig. 2A). We next estimated the latency of troughs and peaks for each ROI and behavioral condition using non-parametric bootstrapping by resampling trials with replacement (1,000 iterations), computing the mean time plots, and extracting the largest negative and positive deflections using *findpeaks* within a −5 to +15 s window. The resulting bootstrap distributions of peak and trough latencies were used to calculate 95% confidence intervals (Fig. 2B).

### Cortical peak-locked data analysis

Global BOLD signal was extracted from the cortical gray matter ROI, defined anatomically with Freesurfer. Power spectra were calculated in 60-s time windows using Morlet wavelets. These epochs were equally divided depending on the average omission rates in that (Extended Data Fig. 2A). The spectra for the drowsy condition were baseline corrected relative to the fully alert condition (Extended Data Fig. 2B). To identify cortical peaks, we filtered the cortical signal in the 0.01–0.05 Hz range and identified the most prominent positive peaks using the *findpeaks* MATLAB function with an amplitude criterion greater than 1 standard deviation of that run (Fig. 3A). The same bandpass filtering was used in the study by Raut and colleagues (2021). We next time-locked both omission likelihood (Fig. 3B) and subcortical fMRI time series (Fig. 3C) to the onset of these cortical peaks (n = 1274). Each time point was then tested for significant change from baseline using hierarchical bootstrapping (1,000 iterations) to account for subject-level variability. Statistical testing was conducted for the −20 s to +20 s window and p-values were corrected for multiple comparisons using the FDR method at the 0.05 level.

We estimated the latency of troughs and peaks for each ROI using non-parametric bootstrapping by resampling trials with replacement (1,000 iterations), computing the mean timeseries, and extracting the largest negative and positive deflections using *findpeaks* within a −25 to +5 s window for the first significant peak/trough and a −15 to +15 s window for the second significant peak/trough. The resulting bootstrap distributions of peak and trough latencies were used to calculate 95% confidence intervals (Fig. 3D). Absolute maximum temporal lags of the subcortical regions relative to the thalamus were calculated from the cross-correlation with maximal lags of - 20 to 20 s. Confidence intervals were calculated using non-parametric bootstrapping by resampling trials with replacement (1,000 iterations), computing the mean of both time series, and calculating the absolute maximum temporal lag for each iteration (Fig. 3E).

### Loss of responsiveness with/without cortical peak data analysis

To further test the hypothesis that the amplitude of AAN regions affects the subsequent global cortical hemodynamic response, we reexamined the loss-of-responsiveness condition (Fig. 2, green traces). We extracted the amplitude values at the maximum of the averaged cortical signal in the loss-of-responsiveness condition (11 s after omission onset, Extended Data Fig. 1A). We then divided trials based on the median of these values, which resulted in two categories: trials with and without a subsequent cortical rise (Fig. 3F). Next, we averaged the fMRI time series for each subcortical region within these two categories and tested for significant differences between them using paired t-tests. The obtained p values underwent multiple-comparison correction using the FDR method at the 0.01 level.

To examine whether subcortical activity during arousal transitions relates to subsequent cortical dynamics, we extended these analyses to account for trial-by-trial variability in cortical peak/trough amplitude. For each time point, fMRI signal changes within subcortical ROIs were tested against baseline using a linear mixed-effects (LME) model of the form: fMRI amplitude ∼ 1 + cortical peak/trough + (1 | subj). This model included a fixed intercept capturing behavior-locked subcortical activity associated with the arousal transition, and a fixed effect of cortical peak/trough amplitude to account for linear variance related to the magnitude of the subsequent global cortical response. Subject-level variability was modeled using random intercepts. The obtained p values for both variables underwent multiple-comparison correction using the FDR method at the 0.01 level (Extended Data Fig. 3).

## Acknowledgments

This research was supported by the AASM Foundation Strategic Research Grant (No. 317-SR-23), the McKnight Scholar Award, the Sloan Fellowship, the Pew Biomedical Scholars Award, the National Institutes of Health awards R01-AG070135, U19-NS128613, U19-NS123717, R21-NS123412, and F31NS139696.

## Data and code availability

All analysis code used in this study will be made available publicly on Github. Processed fMRI data will be shared publicly on manuscript acceptance.

