## Extended Data for "A hypothalamic-brainstem activity sequence underlies arousal fluctuations during daytime drowsiness"

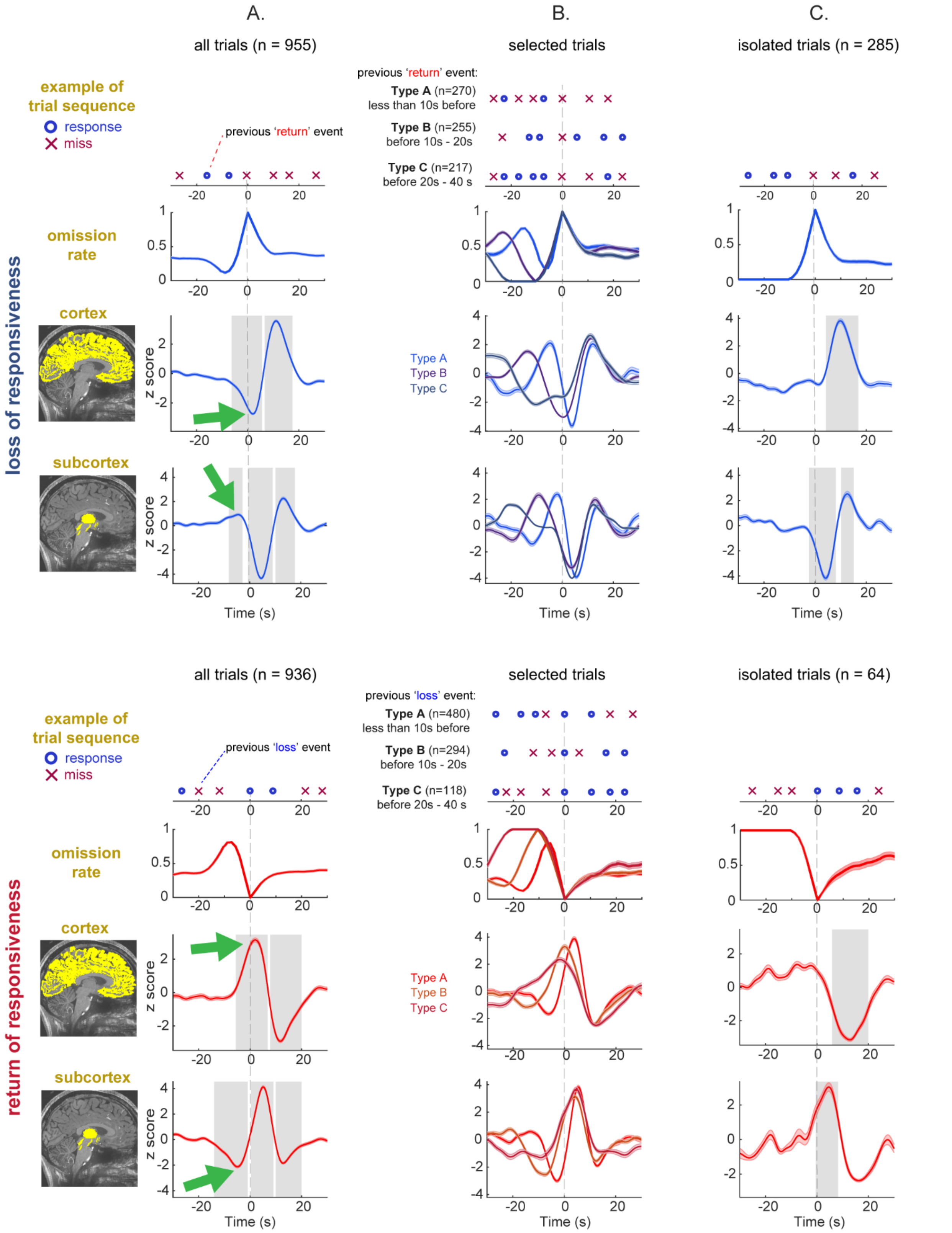


**Extended Data Fig. 1 Event selection method and control analyses for arousal-related hemodynamic responses in global cortical gray matter and combined subcortical regions.** (A) Effect of temporal proximity between arousal transition events when all trials were included (n = 955 and n = 936 for loss- and return-of-responsiveness conditions, respectively). Arousal transition events occur in close temporal proximity (median return–loss interval = 16.2 s; median loss–return interval = 9.8 s), resulting in substantial overlap of preceding events with the event of interest, as revealed by unstable omission-likelihood traces (top row), which may contribute to biphasic signals in both cortex (middle row) and subcortex (bottom row). (B) Control for the influence of preceding arousal events. Trials were subdivided based on the timing of the previous arousal event (type A: <10 s; type B: 10–20 s; type C: 20–40 s before the event of interest), averaged BOLD traces were z-scored, and the timing of peaks and troughs was examined. For both the cortex and thalamus, in both conditions, the first significant peak or trough (see panel A, marked with green arrows; cortex-loss: 2.10 s; cortex-return: 2.26 s; subcortex-loss: −4.51 s; subcortex-return: −4.94 s) showed sensitivity to the timing of the preceding arousal event, whereas the subsequent peaks and troughs were not sensitive, suggesting that they were related to the current arousal event. (C) In line with this conclusion, these early peaks and troughs (see panel A, marked with green arrows) were absent for both the cortex and thalamus in both conditions. Error bars present standard error of the mean. Grey shadings indicate significant changes (linear mixed-effect with random intercept for subject; FDR-corrected $p<0.05$) from the baseline (-30s to -20s).

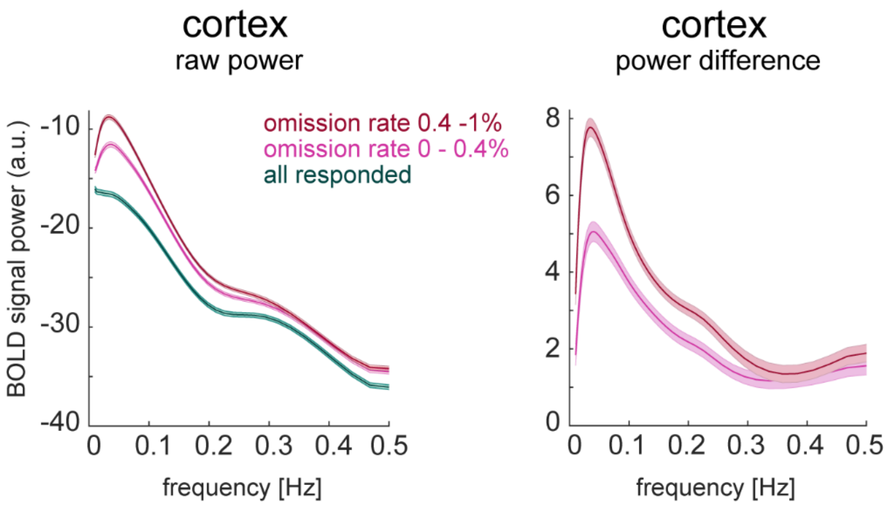


**Extended Data Fig. 2. Spectral characteristics of the global cortical BOLD signal across arousal states**. (A) Global BOLD signal extracted from anatomically defined cortical gray matter. Power spectra were computed for each non-overlapping 60-s epoch using Morlet wavelets and averaged based on omission rates. (B) Power spectra for the two drowsiness levels, baseline-corrected relative to the fully alert condition. The cortical signal showed a broadband power increase during periods of elevated drowsiness, peaking at 0.035 Hz. Error bars present standard error of the mean.

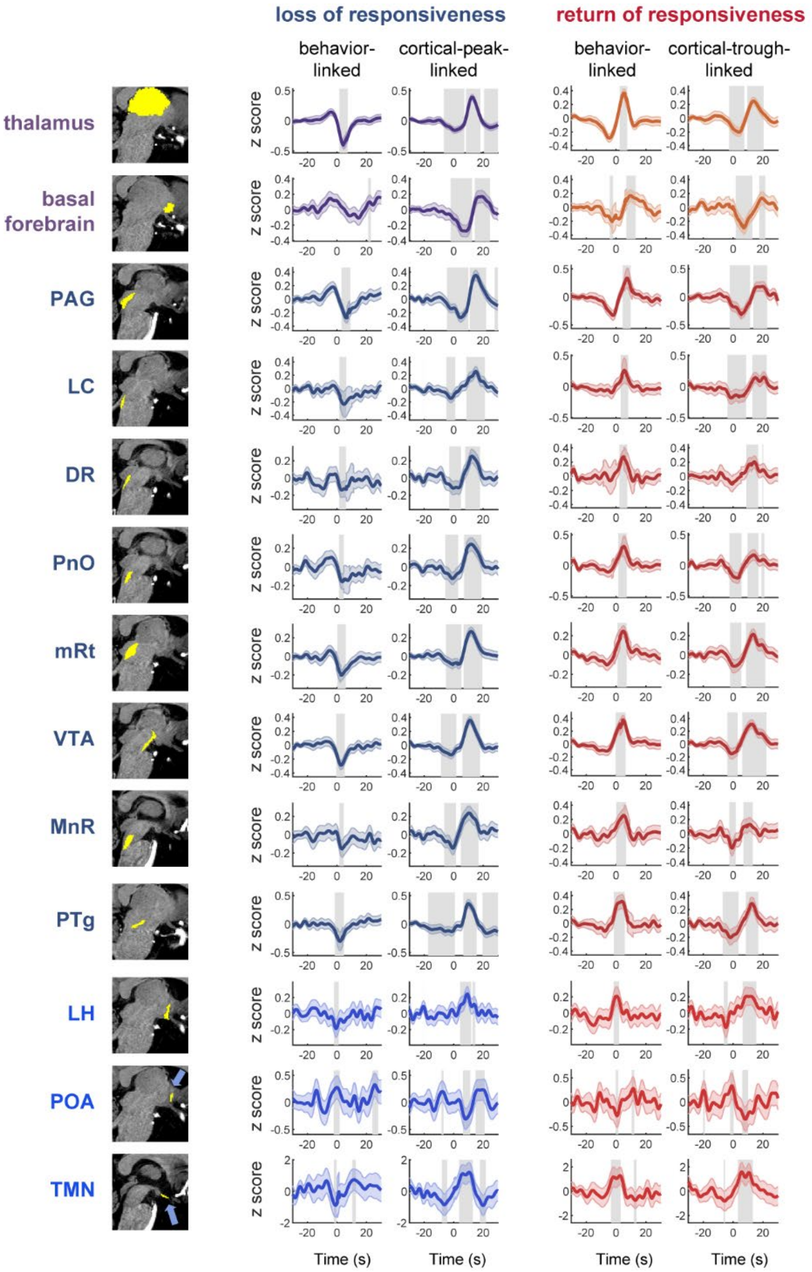


**Extended Data Fig. 3. Behavior-related AAN dynamics are independent of the following cortical response magnitude.**

Results of linear mixed-effects modeling used to test whether arousal-related activity in arousal regulatory system contributes to arousal transitions independently of the magnitude of the subsequent cortical peak (n = 955 and n = 936 for loss- and return-of-responsiveness conditions, respectively). Behavior-locked subcortical activity was captured in the model intercept, while trial-by-trial variability in cortical peak amplitude was included as a separate regressor. Across arousal regulatory regions, behavior-locked dynamics observed during loss and return of responsiveness were preserved after accounting for cortical amplitude variability, indicating independence from the magnitude of the cortical response. Most regions showed a significant relationship with late (11 s after the behavioral arousal event) cortical peak amplitude changes. Interestingly, these effects often occurred earlier than the intercept effect, suggesting network involvement in global vascular responses independent of behavioral responses. Error bars present 95% confidence intervals. Grey shadings indicate significant changes (FDR-corrected $p<0.05$) from the baseline (-30s to -20s).

**Extended Data Table 1 | Latency of subcortical activity changes relative to behavioral state transitions**

| | Loss of responsiveness | | | | Return of responsiveness | | | |
|---|---|---|---|---|---|---|---|---|
| | **Mean (s)** | **Low CI** | **Up CI** | **Type** | **Mean (s)** | **Low CI** | **Up CI** | **Type** |
| Thalamus | 4,4 | 4,2 | 4,7 | trough | 4,9 | 4,7 | 5,2 | peak |
| BF | 8,7 | 6,3 | 9,9 | trough | 8,3 | 7,6 | 9,0 | peak |
| PAG | 6,0 | 5,2 | 6,7 | trough | 6,6 | 6,2 | 6,9 | peak |
| LC | 4,1 | 2,8 | 5,8 | trough | 5,9 | 5,4 | 6,4 | peak |
| DR | 3,2 | 2,6 | 3,7 | trough | 4,7 | 4,3 | 5,1 | peak |
| PnO | 3,3 | 2,8 | 3,9 | trough | 4,2 | 3,7 | 4,7 | peak |
| mRt | 2,8 | 2,5 | 3,2 | trough | 4,4 | 3,9 | 4,9 | peak |
| VTA | 2,8 | 2,4 | 3,2 | trough | 3,8 | 3,1 | 4,4 | peak |
| MnR | 2,6 | 1,9 | 3,2 | trough | 4,1 | 3,1 | 4,7 | peak |
| PTg | 2,4 | 1,8 | 3,0 | trough | 1,7 | 0,8 | 4,1 | peak |
| LH | 0,1 | -0,7 | 1,1 | trough | 0,3 | -0,5 | 1,0 | peak |
| POA | 0,0 | -2,5 | 3,1 | peak | 0,1 | -1,0 | 1,0 | trough |
| TMN | -0,8 | -2,6 | 0,4 | trough | -2,2 | -3,4 | 0,2 | peak |

Latencies (s) to the first prominent peak or trough in fMRI activity relative to the onset of the first omission trial (loss of responsiveness) and the first correct trial following omission onset (return of responsiveness) are shown for each subcortical region of interest (see Fig. 2B). Estimates are derived at the single-trial level, and 95% confidence intervals are obtained using trial-wise bootstrapping.

**Extended Data Table 2 | Latency of subcortical fMRI responses relative to peak of cortical infraslow oscillations.**

| | Early response | | | | Late response | | | | Lag to Thalamus | | | |
|---|---|---|---|---|---|---|---|---|---|---|---|---|
| | **Mean (s)** | **Low CI** | **Up CI** | **Type** | **Mean (s)** | **Low CI** | **Up CI** | **Type** | **Mean (s)** | **Low CI** | **Up CI** | **Type** |
| Cortex | 0,2 | 0,1 | 0,3 | peak | 8,9 | 8,6 | 9,2 | trough | - | - | - | - |
| Thalamus | -7,4 | -7,8 | -7,0 | trough | 1,2 | 1,0 | 1,3 | peak | - | - | - | - |
| BF | -2,4 | -3,0 | -2,0 | trough | 6,0 | 5,3 | 6,8 | peak | 5,2 | 4,8 | 5,5 | positive |
| PAG | -5,3 | -5,7 | -4,9 | trough | 4,0 | 3,6 | 4,6 | peak | 2,8 | 2,6 | 3,0 | positive |
| LC | -7,8 | -9,6 | -5,3 | trough | 2,3 | 1,3 | 4,1 | peak | 1,5 | 1,1 | 2,0 | positive |
| DR | -8,5 | -9,6 | -7,3 | trough | 0,8 | 0,3 | 1,5 | peak | -0,1 | -0,6 | 0,2 | positive |
| PnO | -9,4 | -10,5 | -7,7 | trough | 0,3 | -0,1 | 0,8 | peak | -0,7 | -1,1 | -0,4 | positive |
| mRt | -8,3 | -9,0 | -7,6 | trough | 0,4 | 0,0 | 0,9 | peak | -0,8 | -1,0 | -0,6 | positive |
| VTA | -9,2 | -9,9 | -8,6 | trough | -0,5 | -0,8 | -0,2 | peak | -1,5 | -1,6 | -1,3 | positive |
| MnR | -10,1 | -11,0 | -9,1 | trough | 0,1 | -0,8 | 0,9 | peak | -1,9 | -2,3 | -1,5 | positive |
| PTg | -10,3 | -12,1 | -9,3 | trough | -0,3 | -0,6 | 0,0 | peak | -1,8 | -2,0 | -1,6 | positive |
| LH | - | - | - | - | -0,9 | -2,5 | 0,3 | peak | -2,9 | -3,5 | -2,3 | positive |
| POA | - | - | - | - | -2,7 | -3,7 | -1,5 | trough | -4,0 | -4,5 | -3,5 | negative |
| TMN | - | - | - | - | -2,6 | -4,2 | -1,4 | peak | -5,5 | -6,2 | -4,8 | positive |

Latencies (s) to all significant peaks or troughs in fMRI activity relative to the peak of cortical infraslow waves are shown for each subcortical region of interest (early responses representing initial peaks/troughs and late responses; see Fig. 3D). Maximum time lags (s) between fMRI signals in subcortical regions of interest relative to the fMRI signal in the thalamus are also shown (see Fig. 3E). Estimates are derived at the single-trial level, and 95% confidence intervals are obtained using trial-wise bootstrapping.
